# The biotoxin BMAA promotes dysfunction via distinct mechanisms in neuroblastoma and glioblastoma cells

**DOI:** 10.1101/2022.11.24.517806

**Authors:** Bryan Burton, Kate Collins, Jordan Brooks, Ellie Moore, Karly Marx, Abigail Renner, Kaylei Wilcox, Keith Osowski, Jordan Riley, Jarron Rowe, Matthew Pawlus

**Author notes:** Corresponding author, (MP). These authors contributed equally to this work (Kate Collins, Jordan Brooks, Ellie Moore, Karly Marx, Abigail Renner, Kaylei Wilcox, Keith Osowski, Jordan Riley).

## Abstract

Chronic exposure to the Cyanobacteria biotoxin Beta-methylamino-L-alanine (BMAA) has been associated with development of a sporadic form of ALS called Amyotrophic Lateral Sclerosis/Parkinsonism-Dementia Complex (ALS/PDC), as observed within certain Indigenous populations of Guam and Japan. Studies in primate models and cell culture have supported the association of BMAA with ALS/PDC, yet the pathological mechanisms at play remain incompletely characterized, effectively stalling the development of rationally-designed therapeutics or application of preventative measures for this disease. In this study we demonstrate for the first time that sub-excitotoxic doses of BMAA modulate the canonical Wnt signaling pathway to drive cellular defects in human neuroblastoma cells, suggesting a potential mechanism by which BMAA may promote neurological disease. Further, we demonstrate here that the effects of BMAA can be reversed in cell culture by use of pharmacological modulators of the Wnt pathway, revealing the potential value of targeting this pathway therapeutically. Interestingly, our results suggest the existence of a distinct Wnt-independent mechanism activated by BMAA in glioblastoma cells, highlighting the likelihood that neurological disease may result from the cumulative effects of distinct cell-type specific mechanisms of BMAA toxicity.

## INTRODUCTION

The biotoxin beta-methylamino-L-alanine (BMAA) is produced by several species of cyanobacteria as well as eukaryotic diatoms in both freshwater and marine environments in many locations around the world [1]. The toxin bioaccumulates in the organisms inhabiting local ecosystems and has been identified in Guamanian cycad seed [2]. BMAA has since been detected in the tissues of many other plant and animal organisms inhabiting ecosystems around the world where BMAA-producing microorganisms are located [3]–[6]. Human exposure to BMAA through environmental factors has been implicated in the development of neurodegenerative disorders including amyotrophic lateral sclerosis/ parkinsonism dementia complex (ALS/PDC) in the Indigenous Chamorro population of Guam [7]–[12].

ALS/PDC is a malignant form of ALS found endemic in certain human populations in Guam and the Kii peninsula of Japan [13], [14]. The disorder is caused by widespread neurofibrillary degeneration in the brain and spinal cord of affected individuals resulting in bradykinesis, rigidity, tremor, forgetfulness, and dementia. Currently, no definitive test or treatment exists for the diagnosis or therapy of ALS/PDC. No distinguishing biomarkers have been discovered for the condition and the mechanism by which the disorder develops is fully not understood. Diagnoses are made on the observation of symptoms and the exclusion of other neurological disorders. Therapeutic strategies only address the symptoms of the disorder, not the unknown underlying cause. The onset of ALS/PDC typically begins in a patient’s 40s and symptoms worsen with age. Ultimately individuals progress to a vegetative state and expire 4-6 years after a diagnosis of ALS/PDC [8], [9], [15].

Importantly, symptoms and pathological mechanisms of ALS/PDC bear incredible similarity to other forms of ALS occurring in locations beyond Guam and the Kii peninsula. Studies of ALS/PDC have suggested the possibility that exposure to environmental factors (including BMAA and other biotoxins) could explain the etiology of cases of sporadic ALS and other neurodegenerative disorders around the world [1]. In fact, BMAA has been found in postmortem brain tissue of ALS, Alzheimer’s, and Parkinson’s patients [9], [15], [16].

The toxin BMAA has been demonstrated to exhibit neurotoxicity via excitotoxic mechanisms, however some initial studies rejected the causative relationship of BMAA to ALS/PDC since levels of BMAA found in Chamorro individuals were well below concentrations necessary to produce neurotoxicity [17], [18]. Current evidence strongly suggests that BMAA induces neurodegeneration in primates via a non-excitotoxic mechanism. In these studies, exposure of non-human primates to sub-excitotoxic concentrations of BMAA produced a neurohistopathology similar to that observed in the Chamorro people of Guam. Although these studies have not been repeated in humans due to ethical reasons, they provide strong evidence supporting a causative link between low-concentration BMAA exposure and the development of neurodegenerative disorders in humans via non-excitotoxic mechanisms; a conclusion also supported by other studies in cell culture [10], [19]—[21]. Given that the typical onset of ALS/PDC does not occur until an individual is in their 40s, these results suggest a probable mechanism in which chronic low-concentration exposure to non-excitotoxic doses of BMAA slowly drive neurodegeneration in ALS/PDC patients over several decades.

BMAA has been reported to induce both acute and chronic neurotoxicity via multiple distinct mechanisms [10], [21]—[23]. Acute neurodegeneration has been demonstrated to occur in response to high doses of BMAA in vertebrate subjects[2], [24]. Acute neurotoxicity results through excitotoxic mechanisms dependent upon NMDA and Glutamate receptors. Toxicity via this excitatory mechanism is only observed at exceptionally high doses in vertebrates several orders of magnitude greater than BMAA concentration found in Chamorro subjects with ALS/PDC and is not thought to play a role in the development of this disorder [10], [21], [25].

BMAA is thought to promote neurodegeneration ultimately resulting in ALS/PDC through a non-excitatory mechanism involving altered cell physiology taking place within the nervous systems of organisms exposed to low doses of the toxin over long periods of time. Recent publications have demonstrated that chronic, low-dose exposure to BMAA results in alterations in protein folding, aggregation, expression, enzyme activity, and neuroinflammation, compromising nervous system function at the cellular level [21], [24], [26].

## METHODS

### Cell Culture

IMR-32 cells (ATCC) were cultured in EMEM (Gibco) supplemented with 10% FBS (Gibco) and. U118-MG cells (ATCC) were cultured in DMEM (Gibco) with 10% FBS. Both cell lines were incubated at 37 degrees C and 5% Carbon dioxide.

### Proliferation

MR32 cells were treated with BMAA in a range of different dosages (1, 2, 4 uM) and incubated under normal growth conditions over a period of 48 hours. U118MG cells were treated with BMAA in a range of different dosages (1, 5, 25 uM) and incubated under normal growth conditions over a period of 24 hours or seven days. Cell proliferation was quantified using the XTT proliferation kit (Abnova Cat. No. KA1387) according to manufacturer’s instructions. IMR-32 or U118-MG cells were plated in 96 well plates and treated with BMAA and/or 1 uM XAV939 (Cayman Chemicals Item No. 13596) after 24 hours of incubation at normal conditions.

### Cytotoxcicity

IMR32 cells were treated with BMAA in a range of different dosages (0.8. 1.6, 3.2, 6.4, 12.8, 25 uM) and incubated under normal growth conditions over a period of 72 hours. U118MG cells were treated with BMAA in a range of different dosages (1, 5, 25 uM) and incubated under normal growth conditions over a period of 24 hours or seven days. To detect cell lysis, the presence of extracellular lactate dehydrogenase (LDH) was assessed in cell media at 24, 48, and 72 hours using the CyQUANT LDH Cytotoxicity Assay Kit (Cat. No. C20301) according to the manufacturer’s instructions. Cell lysis was quantified using a Molecular Devices SpectraMax Microplate Spectrophotometer.-

### ROS

IMR32 cells were treated with BMAA in varied doses (1,2, 4 uM) and incubated under normal growth conditions over a period of 48 hours. IMR32 Cells were also treated with 1uM XAV-939 (Cayman Chemicals Item No. 13596), a wnt inhibitor. U118MG cells were treated with BMAA with a single dose (25uM) and incubated under normal growth conditions over a period of 24 hours. U118MG cells were also treated with 10uM EUK-134 an antioxidant (Apexbio Technologies). To detect levels of cellular ROS, an ROS Assay Kit (ABP Biosciences Cat No. A057) was used to quantify DCF fluorescence using a Molecular Devices SpectraMax Microplate Spectrophotometer.

### Neuronal Differentiation

Neuronal differentiation of IMR-32 cells was conducted as previously published. Briefly, cells were plated in 6 well dishes at a density of 50,000/well in normal growth media. After 24 hours of incubation, Cells were treated with BMAA and/or ATRA (ACROS Organics Cat. No. EW-88221-90) at the indicated concentrations by addition to the media and returned to incubation. Cell morphology, analysis of dendrite formation using ImageJ, and RNA extraction for qPCR was conducted following 5 days of incubation post-treatment.

### Plasmids and Transfection

M50 Super 8x TOPFlash was a gift from Randall Moon (Addgene plasmid # 12456; http://n2t.net/addgene:12456; RRID:Addgene_12456); M38 TOP-dGFP was a gift from Randall Moon (Addgene plasmid # 17114; http://n2t.net/addgene:17114; RRID:Addgene_17114); pcDNA-Wnt3A was a gift from Marian Waterman (Addgene plasmid # 35908; http://n2t.net/addgene:35908; RRID:Addgene_35908). Plasmids were ordered from Addgene and were cultured and isolated according to the company’s recommendations. Beta-catenin Activated Reporter assays were conducted using Turbofect Transfection Reagent (ThermoFischer Scientific Cat. No. R0533) to transfect plasmid DNA into IMR-32 cells. Typically, ~50% confluent IMR-32 cells in one well of a 6-well plate were transfected with 200 ng reporter DNA. In addition, 200 ng Wnt3a expression vector was co-transfected to activate the reporter. 24 hours post-transfection, cells were treated by addition of BMAA or CHIR99021(Tocris Cat. No. 2538) to the growth media and incubated under normal growth conditions. 36 hours after transfection, cells were either fixed in 4% PFA and imaged using fluorescent microscopy, collected in RLT buffer for RNA extraction and qPCR, or collected into 400 μl 1x Reporter Lysis Buffer (Promega) and assayed for luciferase activity using a Molecular Devices SpectraMax Microplate Spectrophotometer.

### RNA Isolation & qPCR

RNA was isolated from cells using the RNeasy Plus Micro Kit (Qiagen Cat No. 74034), utilizing DNase to digest possible contaminated genomic DNA. RNA was reverse-transcribed using the SuperScript IV Reverse Transcriptase (Invitrogen Cat. No. 18090010). Levels of mRNA were quantified by SYBR Green qPCR using the AB7500 Real-Time System (AB Biosciences). All primer sets designed to detect target gene mRNA were validated for their product specificity and amplification efficiency using melt curve analysis, qPCR product sequencing, and standard dilution analysis. Amplification efficiencies of primer sets were between 90 and 110%. qPCR results for mRNA were normalized using both GAPDH and beta-actin mRNA. Results were the average of a minimum of three independent experiments performed in triplicate. Primer sequences for mRNA can be found in Table S1.

### Microscopy and Imaging

Cells were observed either in culturing vessels or fixed on slides using an AmScope LB-702 fluorescent microscope under the indicated magnification. Images were captured using the AmScopeAmLite 20200526 microscopy software and analyzed using ImageJ software where appropriate.

### Statistics

Experimental data from a minimum of three replicate experiments for each assay were analyzed by 2-tailed student’s T-test unless otherwise noted and significant p-values reported in each figure; *p<0.05, **p<0.001 vs. untreated cell control; #p<0.05, ##p<0.001 vs. paired experimental sample

## RESULTS

### Neuroblastoma

Neuroblastoma cells have been used extensively to study the effects of BMAA in cell culture [21], [27]. However, the function of BMAA has not been reported in the IMR32 neuroblastoma cell line. Observing the effects of BMAA in IMR32 cells will provide expanded perspective about the toxin’s behavior in a wider range of neuroblastoma cell lines.

### BMAA induces cytotoxicity in IMR-32 human neuroblastoma cells

Acute BMAA exposure has been reported to induce cell apoptosis via excitotoxic mechanisms in neuronal and neuroblastoma cells [27]. By comparison, very little is known about the effects of sub-excitotoxic doses of BMAA on cell signaling events in neuroblastoma cell lines. While BMAA cytotoxicity has been well documented in SH-SY5Y neuroblastoma cells, acute cytotoxicity specifically in the IMR-32 neuroblastoma cell line has not been reported [21], [27]. Treatment of IMR-32 cells with increasing amounts of BMAA revealed an increase in extracellular LDH protein, consistent with increased cell membrane permeability and cytotoxicity (Fig. 1A). Toxicity was observed to increase with both BMAA dose and incubation time and sharply increased at 24, 48, and 72 hour timepoints at BMAA concentrations greater than 4uM. Although Additionally, although minimal cytotoxicity was observed with BMAA doses less than 4uM, changes in cell morphology were observed and characterized by a darkened and rounded, neuroblast-like appearance in contrast to the more elongated, fibroblast-like morphologycells observed in untreated cells (Fig. 1B, quantified in Fig. 1C). The observed cell morphology potentially indicated effects on protein aggregation, cell differentiation or apoptosis.

**FIGURE 1.**
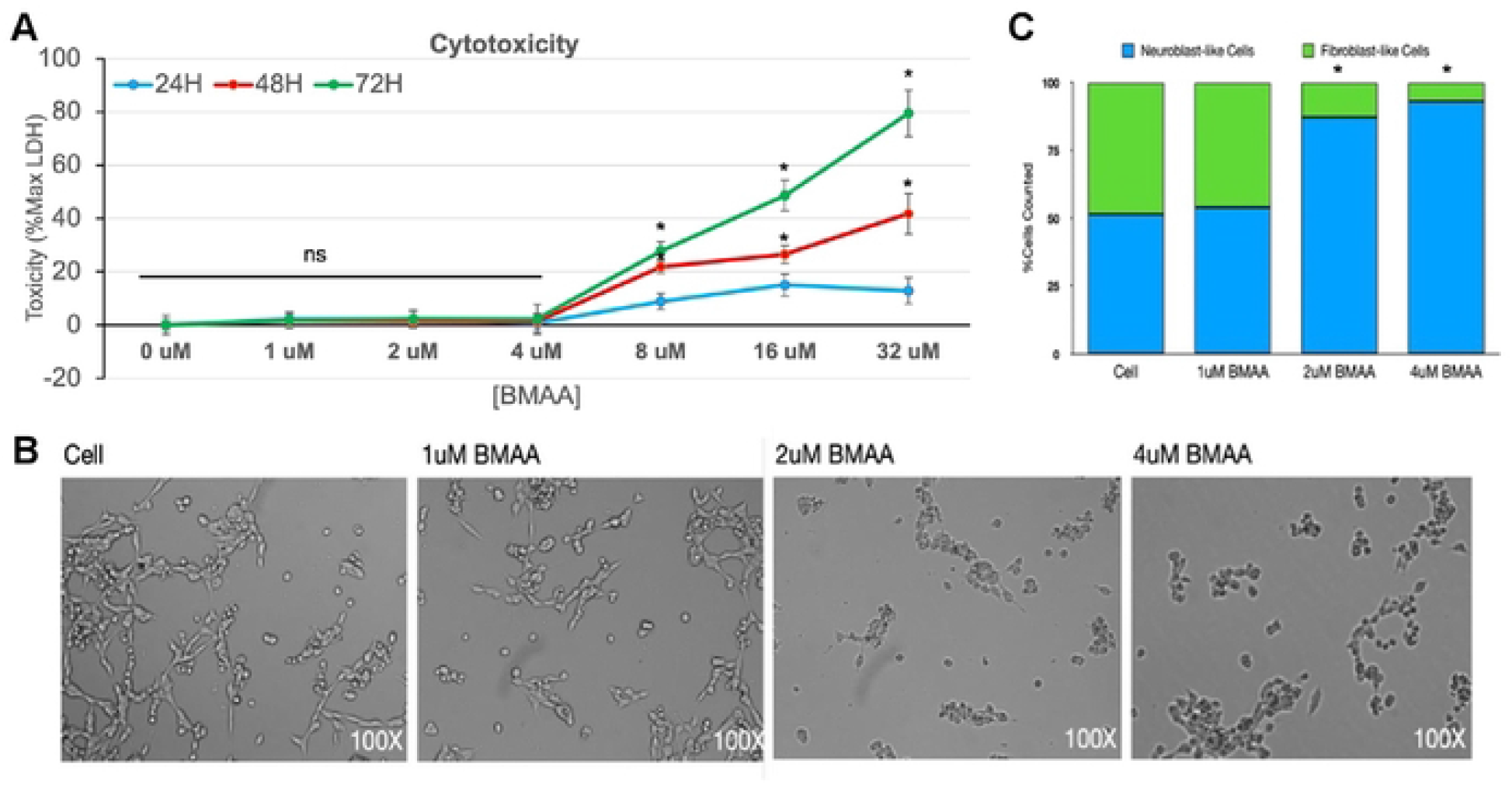
BMAA induces cytotoxicity in IMR-32 human neuroblastoma cells. Cell lysis in response to BMAA was supported by cytotoxicity assay (A). Cells treated with BMAA exhibit morphological changes consistent with cell death (B, quantified in C). Cells treated with several doses of BMAA were incubated under normal growth conditions over a period of 72 hours. The presence of extracellular lactate dehydrogenase (LDH) was assessed in cell media at 24, 48, and 72 hours. The data shown is presented as a percent of maximum LDH (Max LDH) measured from lysed cells. Spontaneous (background) LDH has been subtracted.

### BMAA decreases cell proliferation and increases reactive oxygen species

Because the downstream effects of numerous signaling pathways converge on cell growth effects, we observed the effects of sub-excitotoxic doses of BMAA on proliferation of IMR32 cells in culture (Figure 2). Treatment with increasing doses of BMAA resulted in decreased cell number and inhibition of cell proliferation, as measured by XTT assay (Fig. 2A, B). Although proliferative inhibition trended with increased BMAA dose, statistical significance was not observed at concentrations less than 4uM (Fig. 2B). These findings support the hypothesis that while BMAA may be capable of exhibiting repression of proliferative cells, it may also cause cellular defects via additional processes, especially at low concentrations.

**FIGURE 2.**
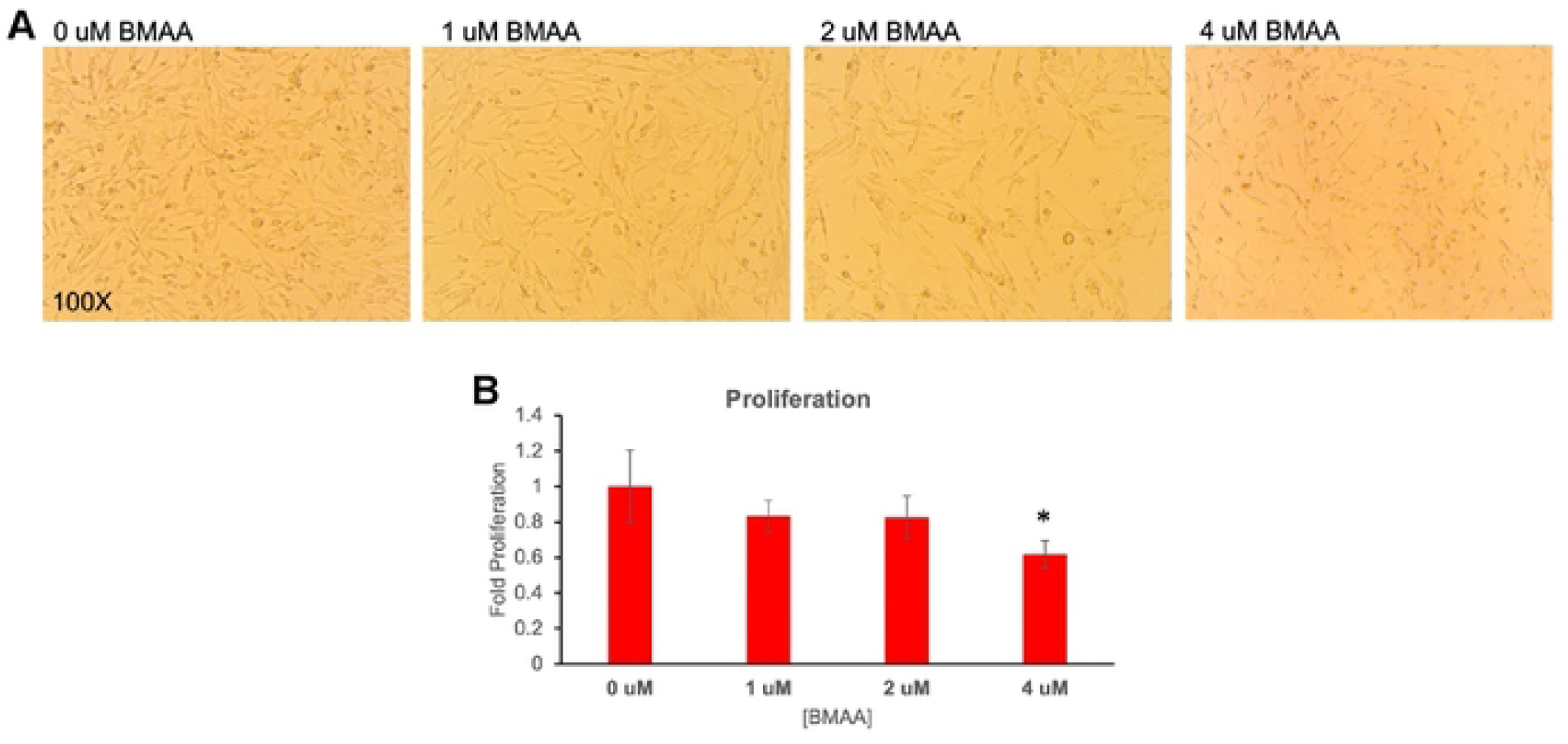
BMAA decreases cell proliferation. Treatment with increasing doses of BMAA resulted in decreased cell number and inhibition of cell proliferation as measured by XTT assay (Fig. 2A, B). Although proliferative inhibition trended with increased BMAA dose, statistical significance was not observed at concentrations less than 4uM (Fig. 2B). These findings support the hypothesis that while BMAA may be capable of exhibiting repression of proliferative cells, it may cause cellular defects via additional processes, especially at low concentrations.

Many neurodegenerative conditions including Alzheimer’s, Parkinson’s, Aging, and ALS are characterized by an increase in cellular accumulation of reactive oxygen or nitrogen species (ROS, RNS) in the brain [13], [28]. In fact, cellular accumulation of ROS or RNS is a known mechanism of toxicity of many neurotoxins [29]urotoxins. ROS accumulation may result from altered cell metabolism, protein aggregation, or organelle damage, and can directly contribute to cellular dysfunction by through damaging lipid membranes, proteins, and DNA damage [29]-[31]. Cellular reactive oxygen species abundance was found to increase in IMR-32 cells exposed to BMAA in a dose-dependent manner in a fluorescence-based cellular assay using the ROS-reactive chemical Dichlorofluorescein (DCF) (Figure 3). Interestingly, although ROS/RNS at high intracellular concentrations contribute to cell damage, dysfunction, and death, ROS at low concentrations have been shown to be important cell signaling molecules and regulate important cell functions in many contexts [32], [33]. These findings suggest that BMAA may exhibit impacts on cell signaling events even at sub-excitotoxic doses by stimulating the accumulation of reactive oxygen species.

**FIGURE 3.**
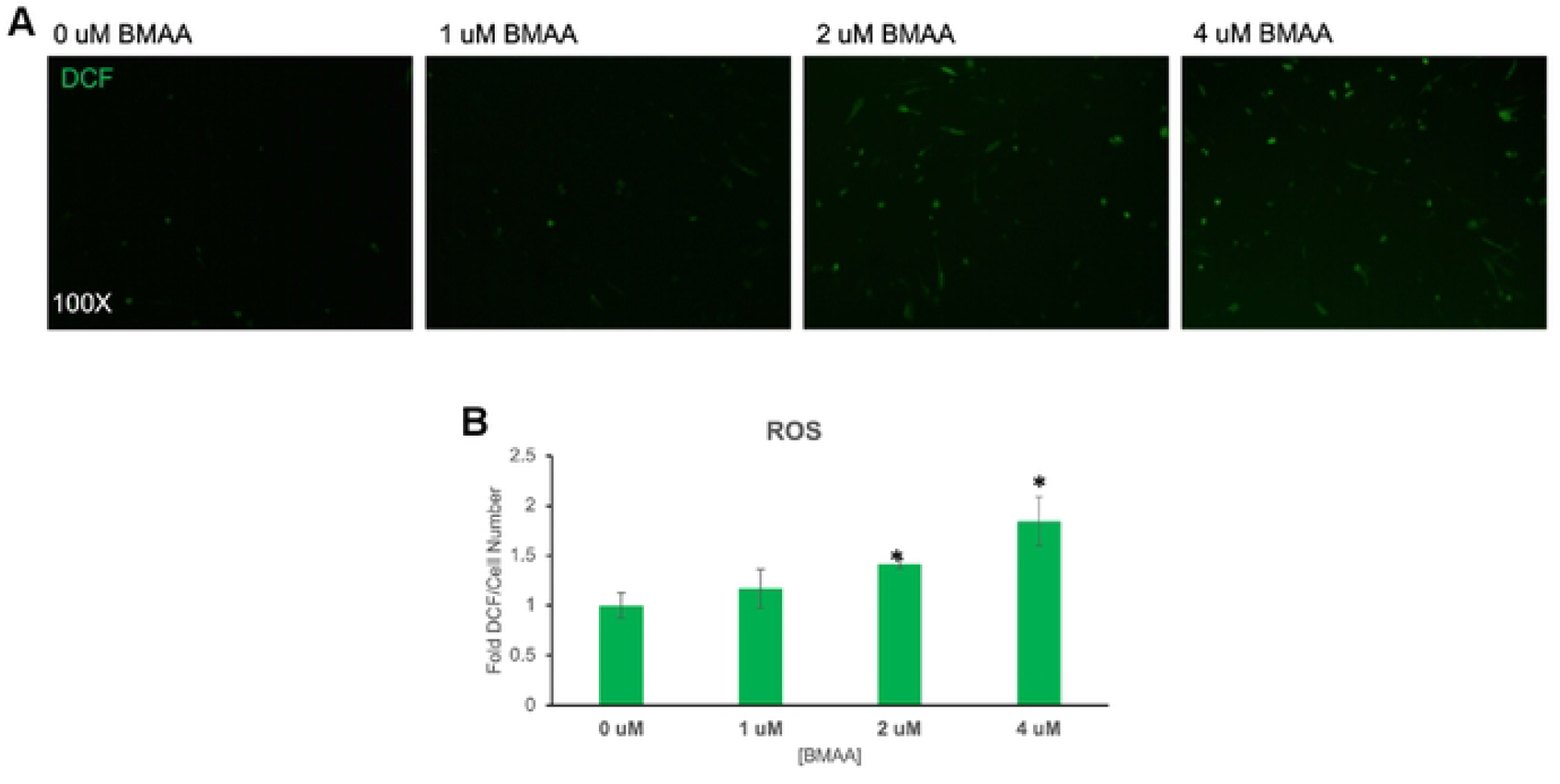
BMAA increases reactive oxygen species. Cellular reactive oxygen species abundance was found to increase in IMR-32 cells exposed to BMAA in a dose-dependent manner in a fluorescence-based cellular assay using the ROS-reactive chemical Dichlorofluorescein (DCF) (Figure 3). Interestingly, although ROS/RNS at high intracellular concentrations contribute to cell damage, dysfunction, and death, ROS at low concentrations have been shown to be important cell signaling molecules and regulate important cell functions in many contexts.

### BMAA activates Wnt/Beta-catenin signaling

The canonical Wnt signaling pathway is critical for appropriate embryonic development and plays many context-dependent roles in cell differentiation and function [34]. Defects in the canonical Wnt signaling pathway have been implicated in many neurodegenerative diseases and neurodevelopmental disorders including AD, PD, ALS, Schizophrenia, and ASD [35]–[37]. Canonical Wnt signaling is reported to regulate cellular proliferation and exhibit significant cross-talk with cellular ROS and other signaling pathways in neural cells [33], [38]-[40]. We wondered if the effects we observed with BMAA treatment could be explained by an induced misregulation of the canonical Wnt signaling pathway. To investigate this possibility, IMR-32 cells were transfected with a Wnt/Beta-catenin activated reporter (BAR), in which GFP or Luciferase were expressed under control of a promoter containing several Beta-catenin response elements. Transfected cells were treated with either recombinant Wnt3a (a ligand of canonical Wnt signaling) or 4uM BMAA. GFP fluorescence and luciferase activity were measured after 48 hours of treatment. Bright-field and fluorescent micrographs were imaged (BF, GFP) (Figure 4A) and GFP fluorescence was quantified using a plate reader. Luciferase chemiluminescent activity was assessed at this time and quantified as well. Both GFP fluorescence and luciferase chemiluminescence were significantly enhanced in cells treated with BMAA, albeit to a lesser extent than in cells treated with rWnt3a (Fig. 4B). The results indicated that BMAA may be capable of inducing canonical Wnt/Beta-catenin activity in treated IMR-32 cells.

**FIGURE 4.**
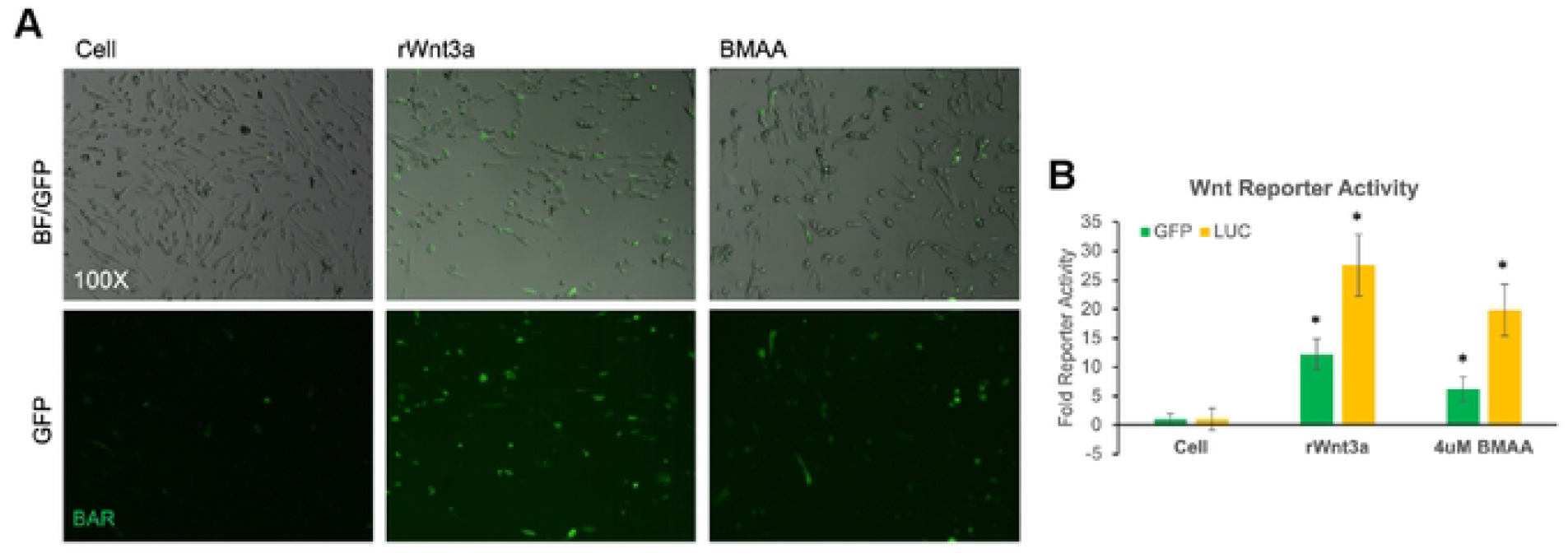
BMAA activates a Wnt/Beta-catenin reporter. IMR-32 cells were transfected with a Wnt/Beta-catenin activated reporter (BAR) in which GFP or Luciferase were expressed under control of a promoter containing several Beta-catenin response elements. Transfected cells were treated with either recombinant Wnt3a (a ligand of canonical Wnt signaling) or 4uM BMAA. GFP fluorescence and luciferase activity were measured after 48 hours of treatment. Brightfield and fluorescent micrographs were imaged (BF, GFP) (A) and GFP fluorescence quantified using a plate reader. Luciferase chemiluminescent activity was assessed at this time and quantified as well. Both GFP fluorescence and luciferase chemiluminescence were significantly enhanced in cells treated with BMAA (B).

In order to further validate the reporter assay results, we next set outsought to determine if BMAA treatment could induce the expression of endogenous Wnt target genes in IMR-32 cells. Wnt target genes are cell-type- and context-dependent[41]. The genes MYC and EPAS1 have both beenare both reported to be targets of canonical Wnt signaling in the neuroblastoma cell line SY5Y and others[42]. We validated the induction of these genes in IMR-32 cells by treatment of cells with recombinant Wnt3a (Figure 5). Interestingly, expression of both MYC and EPAS1 expression was also induced by BMAA treatment as well. XAV939 is an inhibitor of Tankyrase 1 and 2 proteins, whose activity is required for phosphorylation of Beta-catenin and dissociation of the cytoplasmic destruction complex. XAV939 has been conventionally used as an inhibitor of the Wnt/Beta-catenin pathway, as it’s use promotes the cytoplasmic sequestration and degradation of Beta-catenin[43], [44]. We further demonstrated that activation of these Wnt targets by BMAA was likely to be at least partially dependent upon canonical Wnt signaling as XAV939 (XAV), significantly reduced the activation of MYC and EPAS1 by BMAA (Fig. 5, XAV). These results demonstrated that BMAA was capable of inducing endogenous Wnt activity, an integral pathway for and due to the known roles of these genes in cell division and differentiation, and suggested a possible mechanism of BMAA toxicity, in which cell differentiation and development may be affected through modulation of Wnt signaling.

**FIGURE 5.**
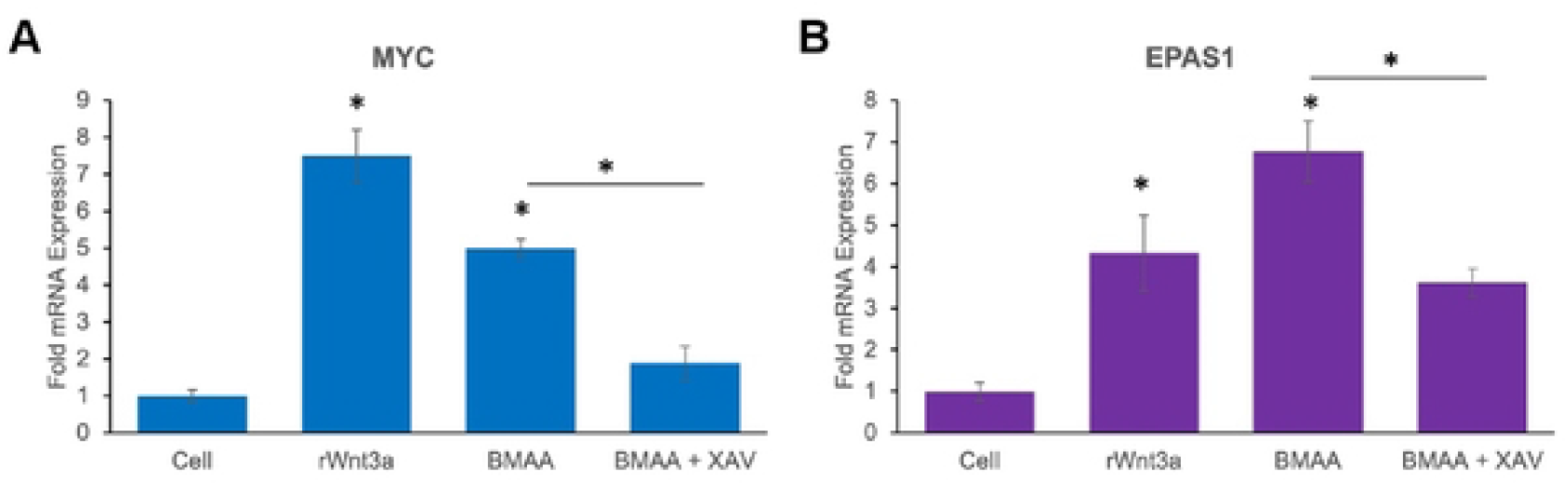
BMAA activates endogenous Wnt target genes. qPCR results demonstrate significant activation of endogenous Wnt target genes MYC (A) and, EPAS1 (B), by 2uM BMAA and a transfected Wnt3a expression plasmid (Wnt3a). Activation of Wnt target genes by BMAA was inhibited by the canonical Wnt pathway inhibitor XAV-939 (XAV). Results indicate average fold expression levels of mRNA relative to Beta actin mRNA expression and represent the average of at least three experiments conducted in triplicate.

### Wnt Inhibition rescues cell function

The canonical Wnt signaling pathway has been demonstrated to regulate myriad cellular processes including proliferation, metabolism, differentiation, and other functions, thiswhich suggested that Wnt inhibition may effectively reverse the functional effects of BMAA on cells [34], [36]. IMR-32 cells co-treated with BMAA and XAV939 exhibited a significant increase in proliferation and decrease in ROS accumulation induced by BMAA (Fig. 2B, 3B; BMAA+XAV). These results indicated that Wnt inhibition may be an effective strategy in reversing some cellular dysfunction induced by BMAA, and is likely mediated through the reduction of endogenous Wnt target genes. Importantly, the finding that BMAA-induced Wnt target gene expression, proliferation, and ROS accumulation could be reversed by a canonical Wnt inhibitor further suggests that these processes occur downstream of Wnt activation by BMAA and are likely to not be responsible for the indirect activation of Wnt themselves.

### BMAA effects neuronal differentiation

Misregulation of Wnt signaling has been well-documented to contribute to neurological disease and carcinogenesis, and can also lead to severe defects in development due its roles in cell differentiation and fate determination[34], [36], [42]. Because Wnt signaling is known to play crucial roles in promoting neuronal differentiation from neural progenitor cells, we next wondered if BMAA might be capable of recapitulating this effect in IMR-32 cells [45], [46]. The neuroblastoma cell line has been used as a cancer cell model of neuronal differentiation, in which treatment with all-trans retinoic acid (ATRA) induces differentiation to neuronal-like cells (Figure 6A). IMR-32 cells were cultured under normal conditions in the presence of ATRA and/or BMAA for a period of 5 days. Cells were then photographed and analyzed to quantify average numbers and length of neurites in all conditions using ImageJ (Fig 6B-D). Interestingly, BMAA treatment during the differentiation protocol resulted in a significantly greater number of neurites, an indicator of neuronal differentiation, especially at a low dose of ATRA (3uM) which was insufficient to efficiently induce neurite formation alone (Fig 6C). Although overall neurite number was found to increase with BMAA treatment, and increase in average neurite length, which can further indicate differentiation, was not observed (Fig 6D). To further verify that neuronal differentiation had been successful, and to evaluate the extent of neuronal differentiation achieved, RNA was extracted from the treated cells and gene expression was analyzed by qPCR. Expression of the neural progenitor marker PAX6 was significantly decreased to a similar level in all cells treated with ATRA, including those co-treated with BMAA, indicating that ATRA treatment successfully pushed cells toward a neuronal lineage. However, eExpression of MAP2, a marker of mature neurons, however was not changed by any treatment, demonstrating that although ATRA treatment may induce differentiation toward neuronal cells, this method does not produce fully mature neurons and this is unaffected by BMAA treatment. Induction of TROY mRNA (a known Wnt target gene [45]) was also observed in cells co-treated with BMAA and ATRA, indicating that BMAA is likely enhancing Wnt activity in treated cells throughout the differentiation process and suggesting that BMAA may enhance neurite formation through Wnt activation. Importantly, while BMAA seems capable of enhancing neuronal differentiation in these experiments it does not appear that BMAA alone is sufficient to induce neuronal differentiation in these experiments, as cells treated with BMAA in the absence of ATRA did not show signs of differentiation.

**FIGURE 6.**
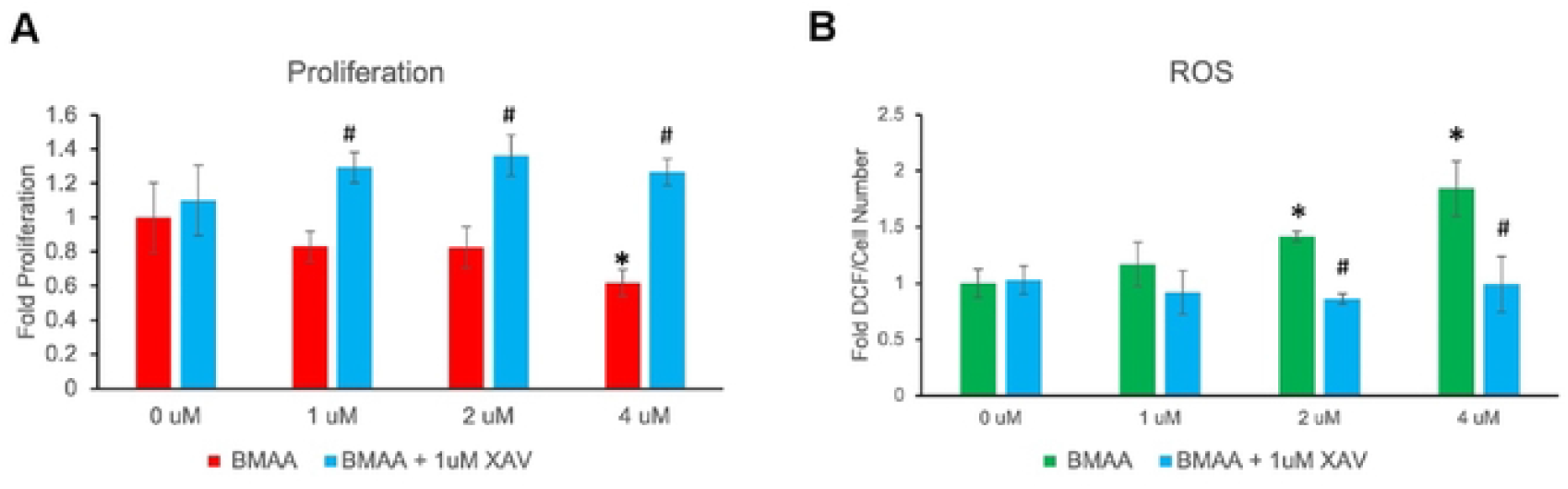
Wnt inhibition rescues cell function. Following treatment with XAV939, an inhibitor of canonical Wnt signaling, BMAA-treated cells were rescued from the proliferation-decreasing effects of BMAA (A). Cellular ROS levels induced by BMAA were also restored by co-treatment with the canonical Wnt signaling inhibitor, XAV-939 (XAV-1uM) (B).

### Glioblastoma

Our findings suggested a mechanism by which sub-excitotoxic BMAA exposure may produce cell dysfunction in proliferation, ROS generation, and differentiation through activation of canonical Wnt signaling in a neuroblastoma cell model. Because the central nervous system is comprised of multiple types of both neuronal and glial cells, and Wnt signaling is known to influence different processes in different cell types, we were curious to characterize the effects of BMAA in a glioblastoma cell model [46].

### BMAA increases glioblastoma cell proliferation

Human U118-MG glioblastoma cells were treated with increasing concentrations of BMAA under normal growth conditions to determine toxicity and effects on cell number. Interestingly, cell number significantly increased with BMAA concentration (Figure 7A). To determine if the observed increase in cell number was due to stimulatory effects on cell proliferation, we performed an XTT assay for proliferative cells (Fig 7B). Results of the assay indicated a significant increase in proliferative cells, however this effect appeared to take time to manifest, as no significant change in cell proliferation was observed at less than 24 hours of treatment (Fig 7B). Interestingly, cytotoxicity was not observed with BMAA in U118-MG cells even at considerably high doses. Cytoplasmic LDH, an indicator of cell lysis, was not increased at doses of BMAA up to 25uM (Fig 7C). Together, these results indicate that BMAA increases glioblastoma cell abundance through activation of proliferative processes and not via inhibition of cytotoxicity. In fact, results of the cytotoxicity assay demonstrate a considerable resistance to BMAA-induced cytotoxicity in these glioblastoma cells.

**FIGURE 7.**
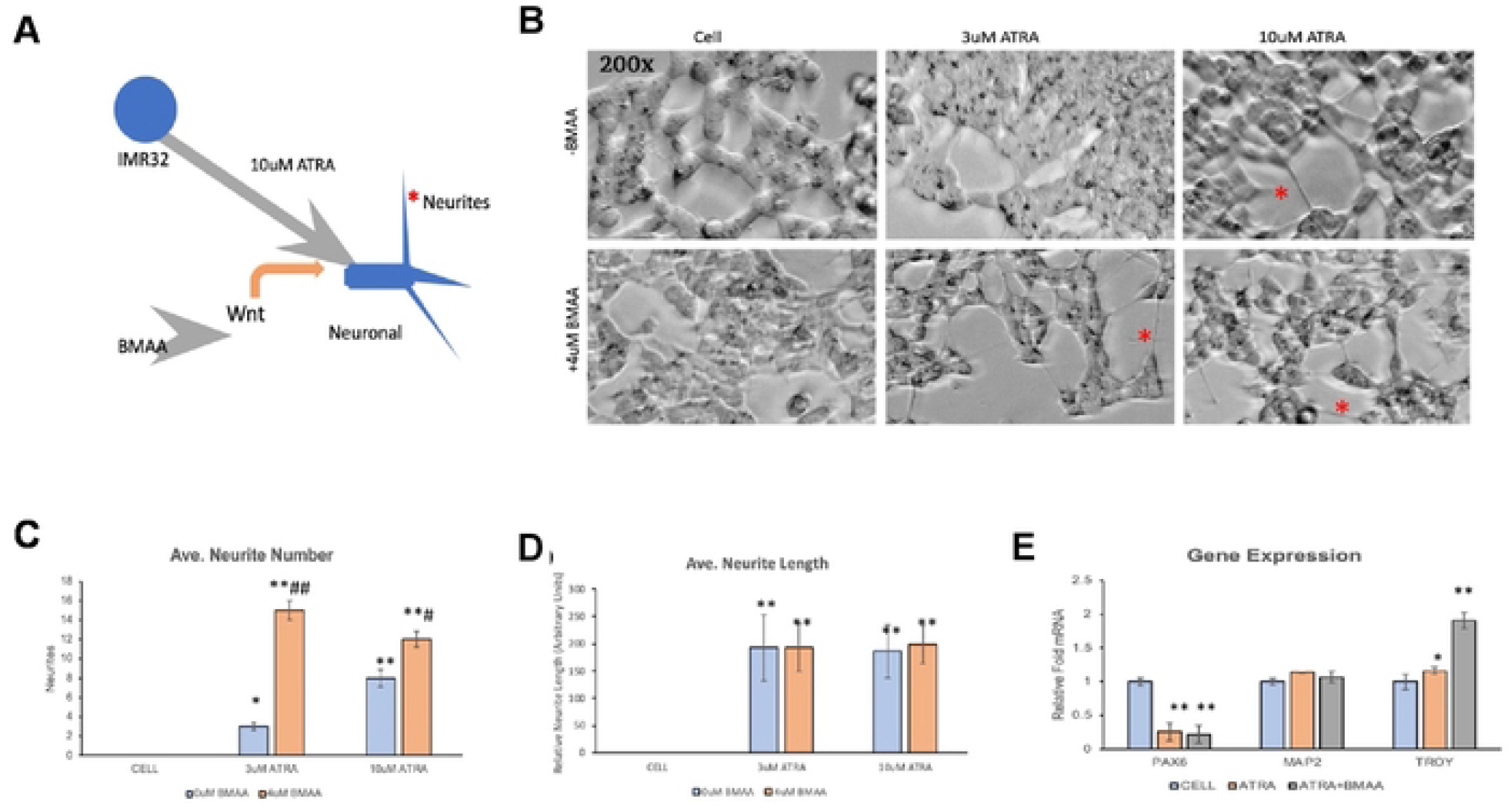
BMAA effects neuronal differentiation. Treatment with all-trans retinoic acid (ATRA) induces neuronal differentiation of IMR32 cells. Wnt enhances this process (A). Micrographs show neuronal differentiation of IMR32 cells treated with BMAA and ATRA for 5 days (B). ImageJ software was used to analyze neurite number and length of 5 images for each treatment (C, D). BMAA increased neurite number even at a lower dose of ATRA, but not length. Gene expression analysis by qPCR showed a decrease in the neural progenitor PAX6 in ATRA treated cells and ATRA+BMAA treated cells showed induction of TROY, a Wnt target and later marker of neuronal differentiation (E).

### ROS are not increased by BMAA in glioblastoma cells

In our neuroblastoma cell model, BMAA induced the accumulation of reactive oxygen species which have the potential to alter cell signaling, damage cell membranes and proteins, and can indicate cellular stress and compromised function[29]—[31]. Although no cytotoxicity was observed in U118-MG cells exposed to BMAA, we hypothesized that BMAA could still be inducing ROS accumulation to some extent and potentially altering proliferative cell signaling pathways. To test this, we performed a cell-based assay for ROS using the fluorescent dye, DCF, in the presence or absence of BMAA (Figure 8). The results indicated that BMAA did not induce ROS accumulation in these glioblastoma cells even when DCF fluorescence was normalized to cell number (Fig 8A, B). The antioxidant compound EUK-134 was used in these experiments as a negative control for ROS and was seen to reduce ROS in all treatments. Interestingly, EUK-134 was also observed to synergize with BMAA in increasing cell number in these experiments, which may suggest that proliferative mechanisms induced by BMAA may be influenced by ROS (Fig 8C, D).

**FIGURE 8.**
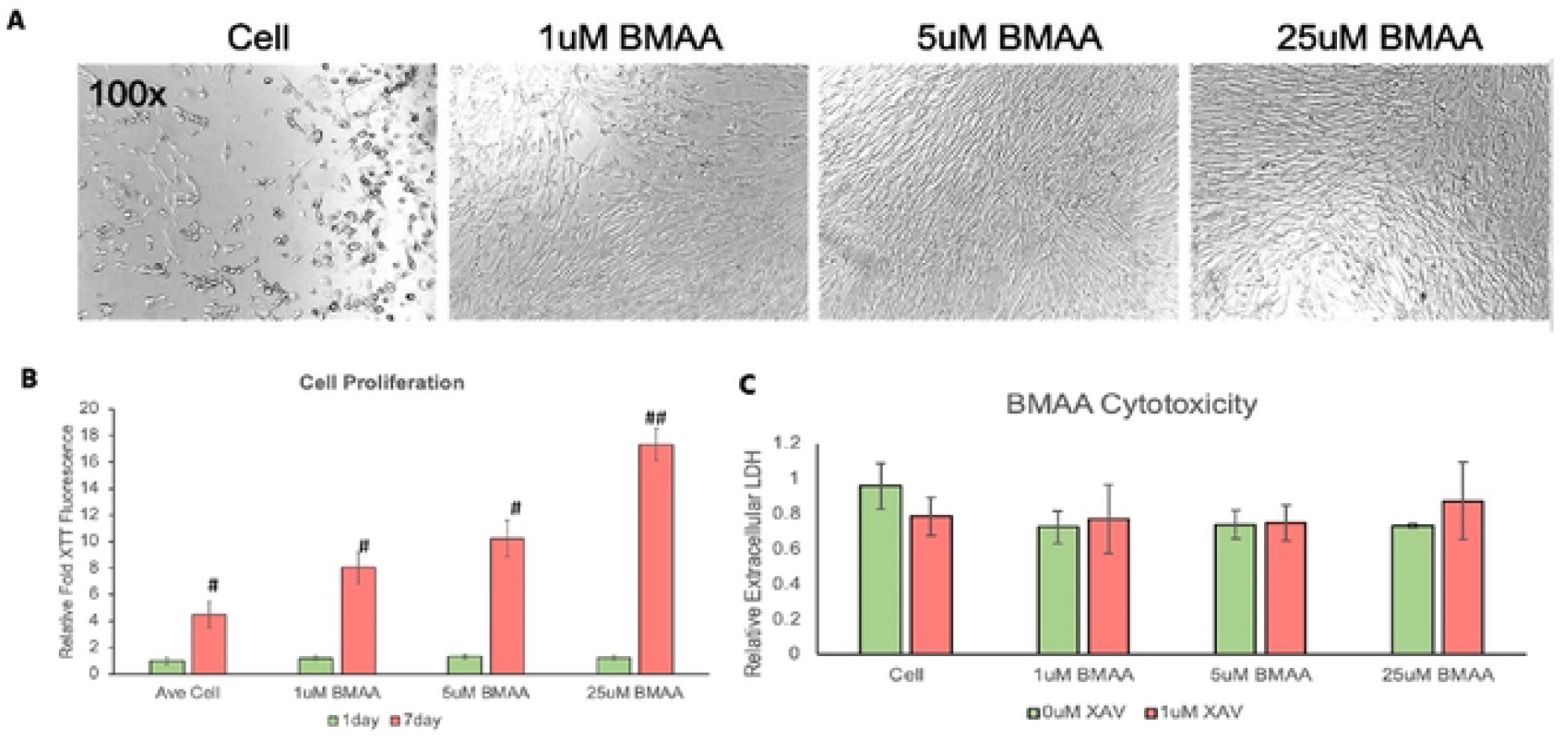
BMAA increases glioblastoma cell proliferation. Cell number is enhanced in the glioblastoma cell line U118MG by BMAA treatment for 7 days (A). XTT assay shows BMAA induces proliferation over a 7 day incubation period (B). BMAA does not increase cytotoxicity in U118MG cells, even at 25uM doses. Cell lysis in response to BMAA was measured by a fluorescence-based LDH cytotoxicity assay (C).

### Wnt signaling is not enhanced by BMAA in glioblastoma cells

Activation of the canonical Wnt signaling pathway has been previously shown to enhance proliferation in human glial cells [47], [48]. Therefore, a general mechanism by which BMAA promotes different functions in different cell types through the activation of Wnt signaling seemed possible. In order to test this idea in glioblastoma cells, we first attempted to transfect U118-MG cells with the Wnt reporter construct we previously used in IMR-32 cells, but these cells were not able to serve as an adequate host for transfection in our hands. We therefore attempted to determine if the Wnt inhibitor XAV939 was capable of reversing the proliferative effects BMAA in these cells (Figure 9). While some level of proliferative inhibition was observed in cells co-treated with BMAA and XAV939, it seems likely that this effect was non-specific for Wnt activity and simply an artifact induced by off-target effects of XAV939, as proliferation was also inhibited in cells treated with XAV939 alone (Fig 9A, B). Gene expression of published Wnt targets was also assessed in U118-MG cells. Interestingly, no induction of the tested Wnt target genes was observed in response to BMAA. In fact, an apparent BMAA dose-dependent inhibition of Wnt target genes was seen (Fig 9C), further suggesting that the effects of XAV939 on cell proliferation were likely due to mechanisms other than decreased Wnt signaling in these cells.

**FIGURE 9.**
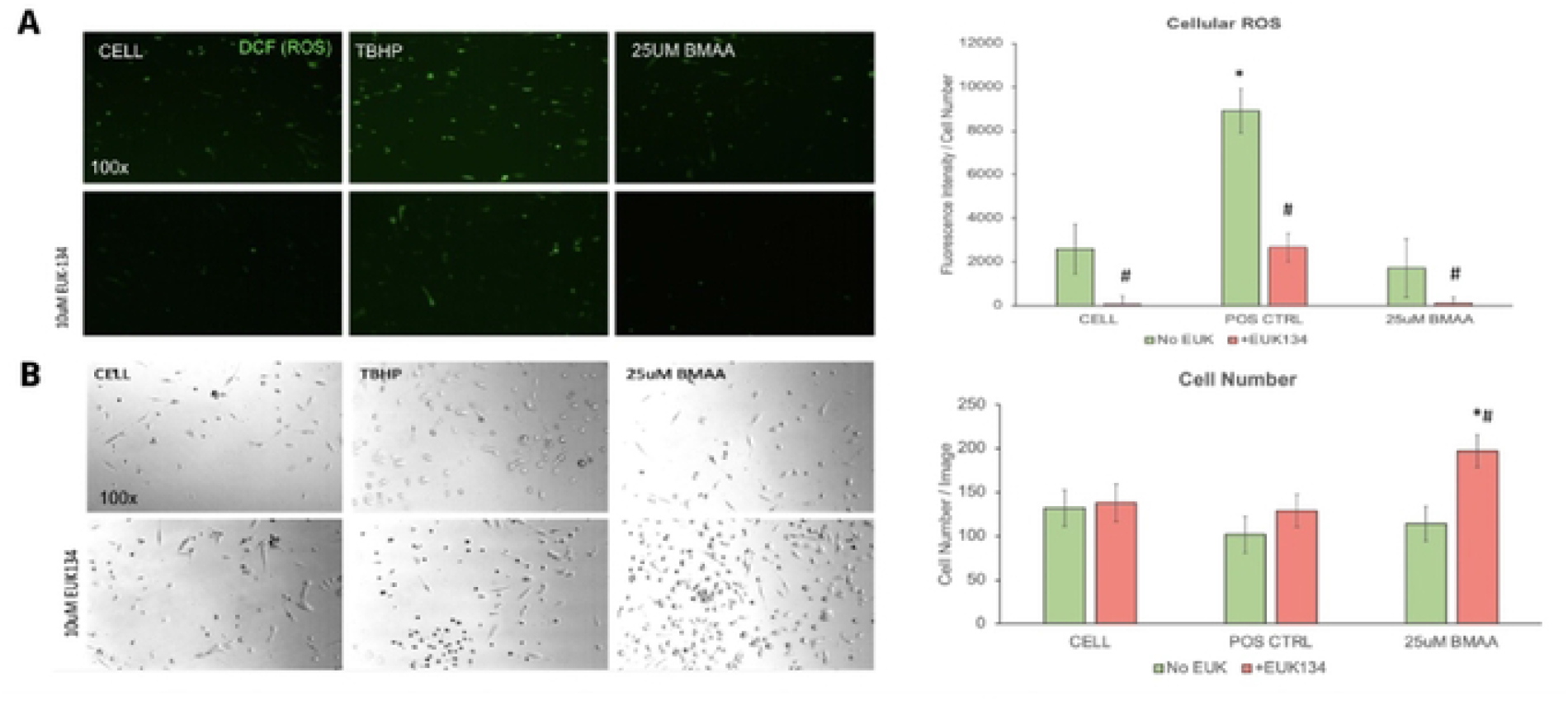
Reactive oxygen species are not increased by BMAA in glioblastoma cells. U118-MG cells treated with BMAA do not show evidence of increased reactive oxygen species following a 24hr exposure to BMAA. Cells subjected to DCF assay for ROS (green fluorescence) are shown in 9A, and ROS-fluorescence normalized to cell number quantified in 9A-right panel. Bright field images demonstrate increased cell number in cells treated with BMAA and the antioxidant EUK-134 (B).

**FIGURE 10.**
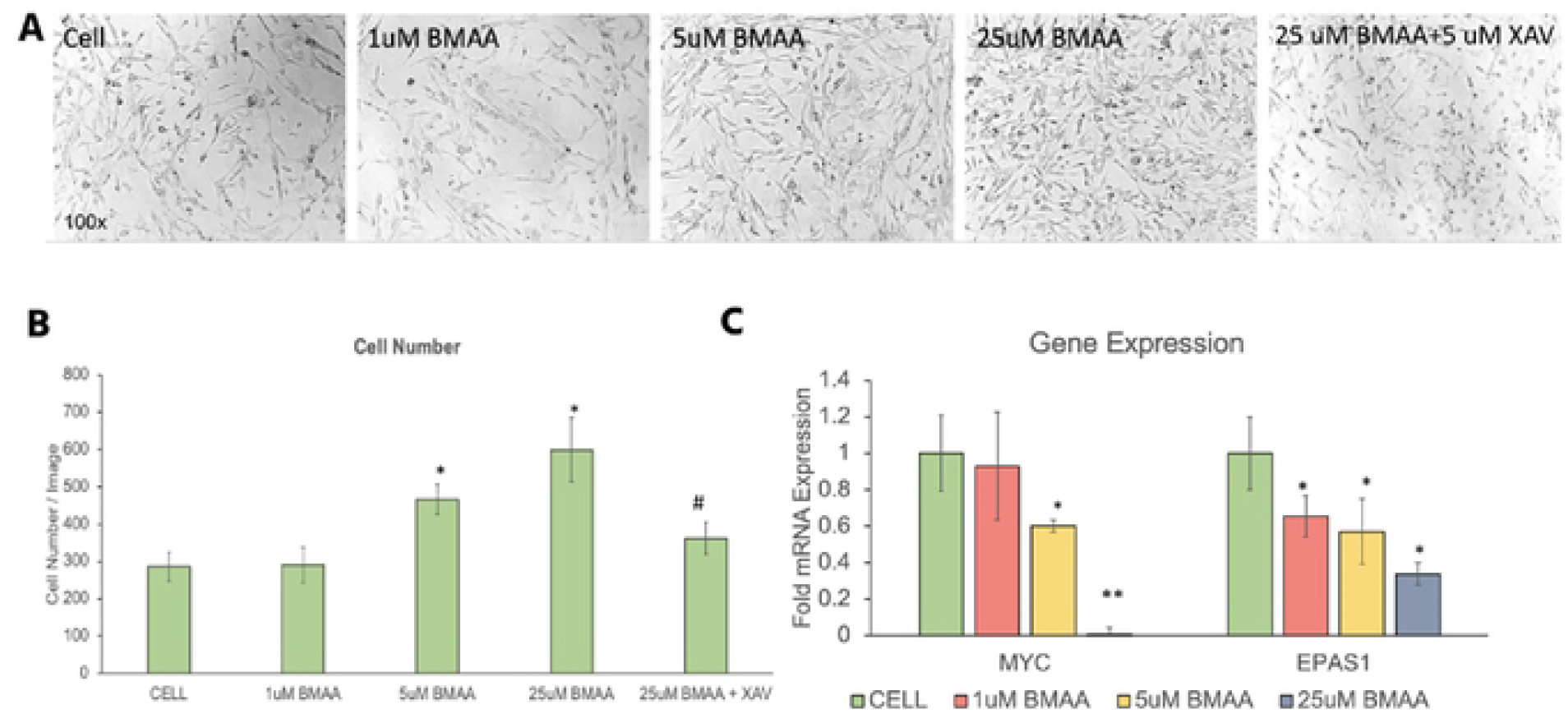
Wnt signaling is not enhanced by BMAA in glioblastoma cells. 48 hour BMAA treatment enhances proliferation and increases cell number in U118-MG cells. This effect is partially inhibited by co-treatment with 1uM XAV-939, a canonical Wnt inhibitor (A, B). Endogenous expression of Wnt target genes, MYC and EPAS1 is not enhanced by treatment with BMAA, suggesting proliferative effects in these cells is not mediated by Wnt pathway up-regulation (C).

### Conclusions and Discussion

Taken together, our data suggest that BMAA produces divergent effects in neuroblastoma and glioblastoma cell lines (Table 1). Interestingly, BMAA appears to promote proliferative defects, ROS accumulation, and effects on differentiation in neuroblastoma cells via a molecular mechanism resulting in activation of canonical Wnt signaling and expression of endogenous Wnt target genes. By comparison, very different effects are observed when glioblastoma cells are treated with BMAA including enhanced proliferation in the absence of ROS accumulation and cytotoxicity. BMAA’s effects in glioblastoma cells appear to be independent of Wnt activation, as Wnt target genes were not induced by BMAA treatment in these cells and an inhibitor of Wnt signaling exhibited minimal effects.

**Table 1:** Effects of BMAA treatment on IMR32 and U118MG cells. BMAA exhibits cell-specific effects in neuroblastoma and glioblastoma cell lines. The toxin inhibits proliferation and enhances cytotoxicity, reactive oxygen species accumulation, neuronal differentiation, and Wnt signaling in IMR-32 neuroblastoma cells. In contrast, BMAA enhances proliferation but does not increase cytotoxicity, reactive oxygen species, or Wnt signaling in U118-MG glioblastoma cells.

| Assay | IMR32<br>(Neuroblastoma) | U118MG (Glioblastoma) |
| --- | --- | --- |
| Proliferation | - | + |
| Cytotoxicity | + | - |
| Reactive Oxygen Species | + | - |
| Neuronal Differentiation | + | N/A |
| Wnt Activation | + | - |

Legend: - did not increase after treatment, + did increase after treatment

Here we provide evidence demonstrating the ability of the cyanobacterial toxin BMAA to induce Wnt signaling in neuroblastoma cells. In the human brain, Wnt signaling has been demonstrated to play critical roles in neural development, stem cell differentiation, regeneration, and function [34], [46]. Due to its involvement in regulating numerous cellular processes, Wnt signaling has been targeted therapeutically for the treatment of many disorders including melanoma, osteoporosis, yet it’s involvement in the etiology of sporadic neurological disorders including ALS/PDC has not been studied [49], [50].

Wnt signaling is known to influence many cellular processes and function in a cell- and contextspecific manner. Importantly ourOur findings therefore suggest that BMAA may promote neurodegeneration and disease via mechanisms involving altered Wnt signaling and downstream effects on numerous biological processes. Especially interesting are our results characterizing BMAA’s apparent effects on cell differentiation. Activation of canonical Wnt signaling has been observed to favor mesodermal differentiation from embryonic stem cells, at the expense of other cell lineages [51]. Wnt has also been demonstrated to influence neural precursor cell self-renewal and enhance neuronal differentiation [52], [53]. In this study we provide evidence that BMAA can enhance endogenous Wnt signaling, and affect neuronal differentiation, suggesting the potential for BMAA to mis-regulate crucial developmental processes. In fact, previous work has demonstrated that treatment of zebrafish embryos with BMAA results in neurodevelopmental defects [54]. These findings may support the idea that while BMAA is capable of inducing acute excitotoxicity at high concentrations, sub-excitotoxic doses of BMAA may promote disease via effects on cell differentiation and development and suggests the prudence of avoiding even miniscule concentrations of BMAA in utero. Indeed, further evidence exists that cellular effects of BMAA may be inherited post-mitotically, again suggesting that a one-time, sub-excitotoxic exposure may be sufficient to promote development of sporadic neurological disease [55].

The reasons that BMAA may exert diverse, cell type-dependent functions in neuronal and glial cells is unclear but is likely to involve cell-specific differences in gene expression, proteome components, and signaling events. One major difference between the cell lines used in this study is the presence or absencearchitecture of NMDA receptors [56]. These receptors are crucial for neuronal cell function and are necessary for excitation and excitotoxicity observed with high concentrations of BMAA. These receptors are found differently expressed inon IMR32 neuroblastoma cells but notversus U118-MG cells. While these receptors (or others) may also facilitate specific cell signaling events driven by sub-excitotoxic BMAA, the differential structure of NMDA receptors on U118-MG cells almost certainly contributes to their resistance to acute BMAA-induced cytotoxicity when compared to IMR32 cells.

Importantly, these results provide evidence that suggest that BMAA induced toxicity and induction of neurodegeneration may be driven by complex interactions between neuronal and glial cells of the central nervous system responding in very different ways to the same toxin and highlight the importance of studying neurodegenerative processes in multiple cell types and complex, interacting, integrated systems. Our findings that BMAA may affect neuronal differentiation further suggest that some sporadic neurodegenerative diseases, including ALS/PDC, may in fact be the end result of mis-regulated developmental or cell differentiation processes. Also suggested is the possibility that other environmental factors associated with the development of neurodegenerative disease may work through similar mechanisms, perhaps involving perturbation of Wnt or other signaling pathways which are crucial for regulating differentiation and development.

## ACKNOWLEDGEMENTS

Research reported in this publication was supported by the South Dakota Biomedical Research Infrastructure Network (SD BRIN) through an Institutional Development Award (IDeA) from the National Institute of General Medical Sciences of the National Institutes of Health under grant number P20GM103443. The content is solely the responsibility of the authors and does not necessarily represent the official views of the National Institutes of Health. This research was performed using WestCore and CCBR Facilities located at Black Hills State University. We thank Mx. Oxana Gorbatenko and Dr. Dan Asunskis for sharing their technical expertise, equipment, and reagents.

---

**TABLE S1.**
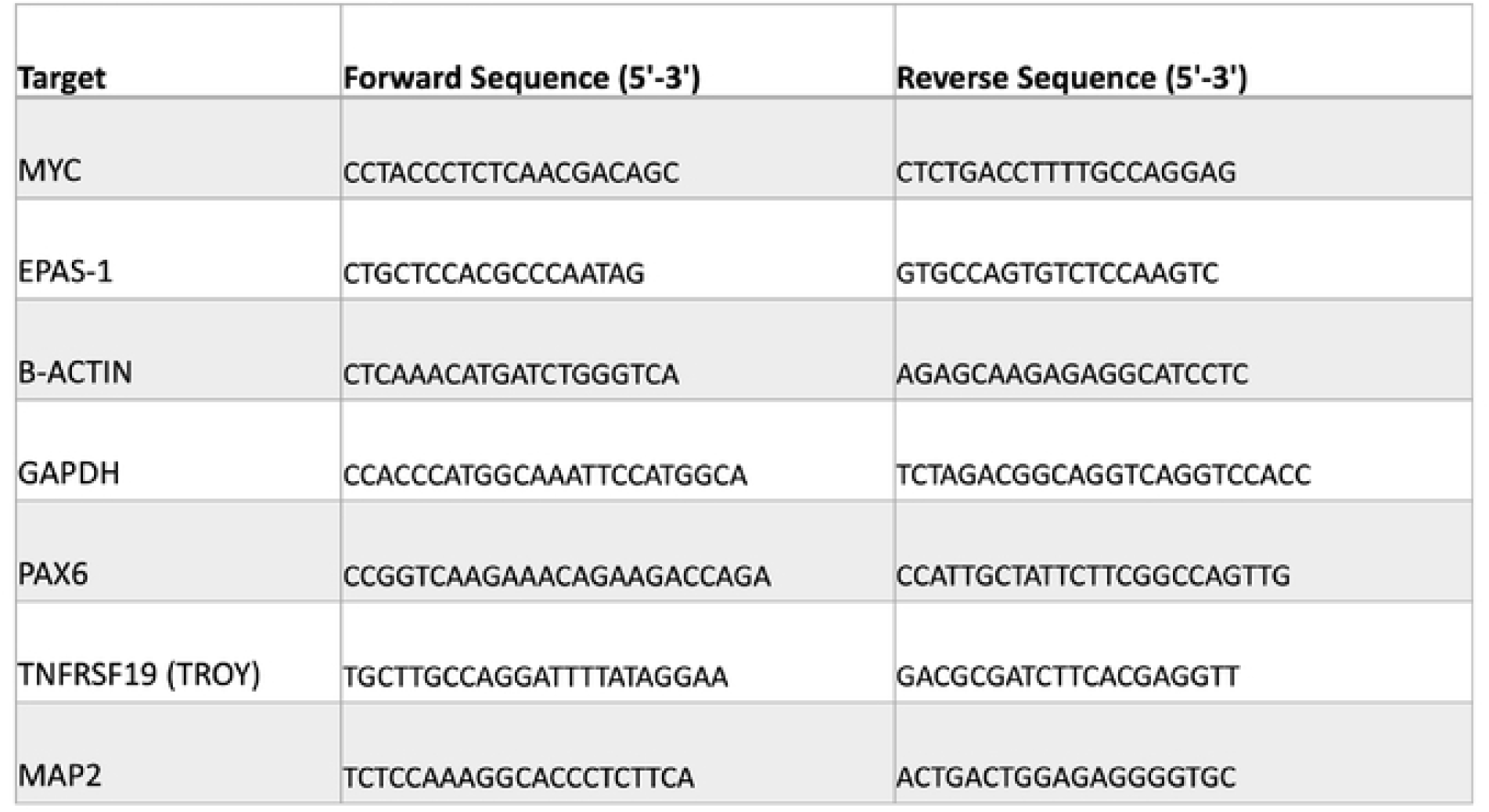
Sequence of qPCR primers used in gene expression analysis.

